# Evolutionary rescue in randomly mating, selfing, and clonal populations

**DOI:** 10.1101/081042

**Authors:** Hildegard Uecker

## Abstract

Severe environmental change can drive a population extinct unless the population adapts in time to the new conditions (“evolutionary rescue”). How does bi-parental sexual reproduction influence the chances of population persistence compared to clonal reproduction or selfing? In this paper, we set up a one-locus two-allele model for adaptation in diploid species, where rescue is contingent on the establishment of the mutant homozygote. Reproduction can occur by random mating, selfing, or clonally. Random mating generates and destroys the rescue mutant; selfing is efficient at generating it but at the same time depletes the heterozygote, which can lead to a low mutant frequency in the standing genetic variation and also affects the establishment probability of the mutation. Due to these antagonistic effects, we find a non-trivial dependence of population survival on the rate of sex/selfing, which is strongly affected by the dominance coefficient of the mutation before and after the environmental change. Importantly, since mating with the wildtype breaks the mutant homozygote up, a slow decay of the wildtype population size can impede rescue in randomly mating populations.

## Introduction

Harsh environmental change can cause a population to decline in size and ultimately go extinct unless adaptation is rapid enough, allowing the population to recover (“evolutionary rescue”). Understanding which genetic and environmental factors influence the evolutionary outcome – population survival or extinction – is of great interest in conservation biology and in the management of drug or pesticide resistance. Over the past few years, several theoretical studies investigated the effect of linkage and recombination between loci on the probability of evolutionary rescue (Schiffers et al., 2013; Bourne et al., 2014; Uecker and Hermisson, 2016). The first two studies use a multilocus model to describe adaptation to climate change, observing that linkage slows adaptation down. The third study, based on a generic two-locus model, finds that a monotonic increase or decrease as well as a maximum or minimum in the probability of rescue as a function of recombination are possible, where the pattern depends on epistasis before and after the environmental change as well as on pronounced stochastic effects. While recombination is an important component of sexual reproduction, the role of sex and the role of recombination in rescue are non-identical problems. For example, sexual reproduction also includes segregation, and recombination happens during other modes of reproduction as well (cf. viruses or homologous recombination in bacteria). Moreover, sexual reproduction can be bi-parental or occur by selfing, adding yet another aspect. The form of reproduction is a potentially important factor for the survival chances of a population.

Evolution has brought about an astonishing variety of reproductive strategies, broadly classified as sexual and asexual. Hermaphrodites are often able to self-fertilize while other hermaphroditic species have evolved refined self-incompatibility systems. A huge body of research has been devoted to disentangling the costs and benefits of the various forms of reproduction, focusing on the rate of adaptation (traditionally in populations of constant size), the colonization of new habitats, the accumulation of deleterious mutations, and the maintenance of standing genetic variation (for review articles see e.g. Stebbins, 1957; Barton and Charlesworth, 1998; Otto, 2009; Hartfield and Keightley, 2012; Barrett, 2013). Bi-parental sexual reproduction is often said to be beneficial for adaptation since it generates genetic variation, giving natural selection the chance to become active. However, neither is it true that sexual reproduction always increases variation nor is the generation of variation necessarily beneficial (Otto, 2009). Generally, sexual reproduction takes a double-edged role in adaptation since segregation and recombination can efficiently create favorable gene combinations (which would need to arise by sequential mutation events in clonal populations) but the very same processes can also break up well-adapted gene complexes. Despite the significance of the topic, little work has explicitly addressed the question of how the mode of reproduction affects the capability of populations to cope with severe environmental change. In their review on studies of evolutionary rescue, Carlson et al. (2014) list a single experimental study for the role of sexual reproduction in rescue. This study, Lachapelle and Bell (2012), finds that sexual reproduction increases the survival chances of the algae *Chlamydomonas reinhardtii* as salt concentration is increased. Likewise, Morran et al. (2009) show that obligate outcrossing populations of *Caenorhabditis elegans* adapt more easily to harsh environmental change than obligate selfing populations. For theoretical work on the role of sexual reproduction in rescue (beyond of recombination), the table in Carlson et al. (2014) shows a gap. Gl´emin and Ronfort (2013) – not cited in the review – analyze a diploid one-locus model for evolutionary rescue in partially selfing populations. However, Gl´emin and Ronfort (2013) exclusively focus on scenarios in which the heterozygotes have a fitness considerably larger than one. In this case, establishment of the heterozygote is sufficient for rescue and the positive or negative effect of segregation in a randomly mating population (generation or destruction of the mutant homozygote) is negligible.

A better understanding of the link between the reproductive strategy and extinction risk is of great interest both in order to understand why we see this variety and distribution of reproductive modes and to predict the impact of current anthropogenic change on species’ diversity. It is of equally great importance for the development of successful strategies to prevent the emergence of resistance in pathogens, insects, or weeds. We need to manage resistance in a broad spectrum of organisms that vary greatly in their mode of reproduction, including bi-parental sexual reproduction (e.g., *Anopheles*, the grass weed *Lolium rigidum*), clonal reproduction in diploids (e.g., the wheat rust fungus *Puccinia triticina*) and haploids (e.g., *Salmonella*), selfing (e.g., horseweed), and combinations of these three modes (e.g., pea aphids, *C. elegans*, *Candida albicans*, *Plasmodium*). Knowing how the mode of reproduction affects the emergence of resistance is crucial for choosing the best treatment plan. For example, different management strategies might be required for weeds that are primarily selfing versus weeds that are primarily outcrossing.

For this article, we compare rescue in randomly mating, selfing, and clonal populations and consider how the rate of sex and the rate of selfing affect evolutionary rescue. We focus on the simplest mechanistic aspect of adaptation in which the three modes of reproduction differ, i.e. on the generation and establishment of homozygote mutants. As in Glémin and Ronfort (2013), there is a discrete change in the environment and adaptation to the novel environment depends on one locus; for most part, we assume that only the mutant homozygote has fitness greater than 1 in the new environment. For clonal populations, we include the possibility of mitotic recombination as an additional pathway to rescue beyond two-step mutation. Adaptation to the novel environment can happen from standing genetic variation or de-novo. There are hence two phases – before and after the environmental change – during which the mode of reproduction influences the probability of population survival by shaping the genetic composition of the population.

After introducing the model, we briefly discuss the advantages and disadvantages of the three modes of reproduction. Subsequently, we provide a more detailed analysis of the effect of the rate of sex and selfing on rescue in separate sections. We derive analytical approximations for the probability of evolutionary rescue, complemented by computer simulations, in order to develop an intuitive understanding of the main principles underlying rescue in randomly mating, selfing, and clonal populations. In the main text, we stick to simple intuitive approximations. A more careful analysis can be found in the supporting information.

## The model

We consider a panmictic population of diploid individuals with non-overlapping generations that is exposed to a sudden severe change in its environment. As a consequence, the population size *N* = *N*(*t*) declines, and the population risks extinction. Adaptation to the new conditions relies on an allelic change at a single locus. Focusing on this locus, we distinguish three genotypes: the wildtype *aa*, the mutant heterozygote *Aa*, and the mutant homozygote *AA*. The numbers of the three possible genotypes are denoted by *n*_*aa*_, *n*_*Aa*_, and *n*_*AA*_, respectively, and *N* = *n*_*aa*_ + *n*_*Aa*_ + *n*_*AA*_. Prior to the environmental change, the population is well-adapted and constant at size *N*_0_. The mutant allele is deleterious under these conditions, and the three genotypes *aa*, *Aa*, and *AA* have relative fitnesses 1, 1 + *δ*_*Aa*_, and 1 + *δ*_*AA*_ with *δ*_*Aa*_, *δ*_*AA*_ < 0. We are most interested in scenarios in which the mutant allele is only present at low frequency at mutation-selection balance. After the environmental change, the absolute fitness of the wildtype drops to 1 + *s*_*aa*_ < 1 such that the wildtype population size declines exponentially. The mutant heterozygote and homozygote have absolute fitnesses 1 + *s*_*Aa*_ and 1 + *s*_*AA*_, and type *AA* is the fittest type in the population. We mainly focus on *s*_*Aa*_ < 0, *s*_*AA*_ > 0, i.e., only the mutant homozygote can grow in the new environment. The heterozygote can be fitter than the wildtype (*s*_*Aa*_ > *s*_*aa*_) or less fit (*s*_*Aa*_ ≤ *s*_*aa*_) such that the population needs to cross a fitness valley. We assume a hard carrying capacity *N*_0_. If *N* > *N*_0_, *N* − *N*_0_ individuals are randomly removed from the population. With fewer than *N*_0_ individuals, fitness is density independent.

We compare three modes of reproduction – clonal reproduction, selfing, and random mating and consider partially clonal and partially selfing populations. Depending on context, we refer to random mating as sexual reproduction or outcrossing. For partially clonal populations, individuals reproduce by random mating (sexually) with probability *σ*_sex_ and clonally (asexually) with probability 1 − *σ*_sex_. For partially selfing populations, individuals reproduce by selfing with probability *σ*_self_ and by random mating (outcrossing) with probability 1−*σ*_self_.

We assume that *σ*_sex_ and *σ*_self_ are independent of population density. We furthermore assume that sexually reproducing individuals produce so many gametes that fertilization is guaranteed; selection acts via viability not via fertility. During reproduction, mutations happen with probability *u* in both directions.

A formal description of the model with the precise life cycle can be found in SI section S1. In the simulations, we let the population evolve for a large number of generations before the environment changes (increasing the number of generations did not affect the results). We subsequently follow the population until it has gone extinct or until the rescue type *AA* has reached 20% carrying capacity. The simulation code written in the *C* programming language makes use of the *Gnu Scientific library* (Galassi et al., 2009).

## Analysis and Results

The probability of rescue is determined by the genetic composition of the standing genetic variation, the efficiency with which the rescue type is generated de-novo, and on its establishment probability. Roughly, this can be summarized in the equation

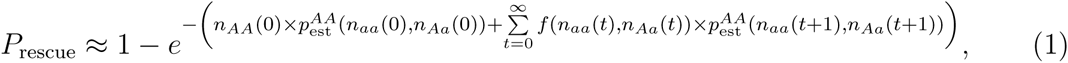

where *n*_*AA*_(0) is the number of mutant homozygotes at the time of environmental change, the function *f* captures the generation of mutant homozygotes in the new environment, and 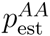 is the establishment probability of type *AA* once generated. The approximation is the Poisson probability for having at least one rescue event and is based on the assumption that rescue mutants reproduce independently from each other as long as they are rare. Unless reproduction is purely clonal or purely selfing, both *f* and 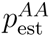 depend on the genotypic composition of the population. We ignore dependence of *f* and 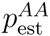 on *n*_*AA*_ in the equation, since the rescue type *AA* is rare in the early phase when the fate of the population is decided (or there is no risk of extinction). We apply branching process theory to estimate the establishment probability 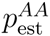. In order to account for fluctuations in *n*_*AA*_(0), we need to average Eq. (1) over the distribution of mutant homozygotes in the standing genetic variation. To do so, we derive an approximation for the corresponding probability generating function in the supplementary material. With this approach, we analyse a set of limiting cases that we discuss in detail below. The detailed analysis can be found in the supplementary material.

### The advantages and disadvantages of the three modes of reproduction

Table 1 summarizes how the mode of reproduction affects the genotypic composition of the standing genetic variation, the rate of de-novo generation of the mutant genotypes, and the establishment of the rescue type.

**Table 1:** The three determinants of rescue. The table summarizes how clonal, selfing, and randomly mating populations differ in the composition of the standing genetic variation, the de-novo generation of the rescue genotype, and in its establishment probability (see text for details). For the approximations of the expected number of heterozygotes and homozygotes in the standing genetic variation, 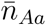 and 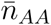, see SI section S3, (S5.3) and (S5.7), (S6.2) and (S6.4).

| Clonal | Selfing | Random mating |
| --- | --- | --- |
| **standing genetic variation** | **standing genetic variation** | **standing genetic variation** |
| many heterozygotes<br>$\bar{n}_{Aa} \approx -\frac{2uN_0}{\delta_{Aa}}$<br>homozygotes:<br>fitness-dependent<br>$\bar{n}_{AA} \approx \frac{2u^2N_0}{\delta_{Aa}\delta_{AA}}$ | few heterozygotes<br>$\bar{n}_{Aa} \approx 4uN_0$<br>homozygotes:<br>fitness-dependent<br>$\bar{n}_{AA} \approx -\frac{uN_0}{\delta_{AA}}$ | many heterozygotes<br>$\bar{n}_{Aa} \approx -\frac{2uN_0}{\delta_{Aa}}$<br>few homozygotes<br>(≈ independent of fitness)<br>$\bar{n}_{AA} \approx \frac{u^2N_0}{\delta_{Aa}^2}$ |
| **de-novo generation** | **de-novo generation** | **de-novo generation** |
| heterozygotes<br>$2un_{aa}(t)$<br>homozygotes:<br>inefficient<br>$un_{Aa}(t)$ | heterozygotes<br>$2un_{aa}(t)$<br>homozygotes: very efficient<br>(but loss of heterozygotes)<br>$\frac{1}{4}n_{Aa}(t)$ | heterozygotes<br>$2un_{aa}(t)$<br>homozygotes: depends on<br>the genotypic composition<br>$\frac{1}{4} \frac{n_{Aa}(t)^2}{N(t)}$ |
| **establishment** | **establishment** | **establishment** |
| fitness-dependent<br>$\approx 2s_{AA}$ | fitness-dependent<br>$\approx 2s_{AA}$ | fitness-dependent<br>depends on genotypic composition<br>$p_{\text{est}}^{AA}(n_{aa}(t), n_{Aa}(t))$ |

For all three modes of reproduction, mutant heterozygotes are generated from wildtype homozygotes by mutation and hence arise at rate *∼* 2*un*_*Aa*_. The formation of mutant homozygotes, however, differs. In clonal populations, a second mutation is required, and the rescue type is generated at a very low rate *∼ un*_*Aa*_ (for mitotic recombination, see below). In fully selfing populations, one quarter of all offspring of a mutant heterozygote are mutant homozygotes. The latter hence arise at rate 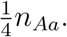. In randomly mating populations, mutant homozygotes can also be generated by segregation and union of mutant gametes. Unlike in selfing populations, the rate depends, however, on the genotypic composition of the population 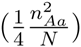. The efficient generation of the rescue type in selfing populations goes hand in hand with the depletion of heterozygotes. The number of heterozygotes is hence much lower in selfing than in randomly mating or clonal populations. Finally, the rescue type does not only have to be generated, it also needs to increase in frequency. In clonal and selfing populations, this only depends on the fitness of type *AA*. In randomly mating populations, mating between the two homozygote types breaks the rescue type up. Its “effective selection coefficient” is reduced from *δ*_*AA*_ and *s*_*AA*_ to roughly 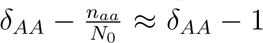 and 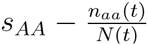, respectively. This has two consequences. While in clonal and selfing populations, the frequency of the mutant homozygote *AA* prior to the environmental change strongly depends on its fitness, the rescue type remains at a low frequency in randomly mating populations before the environmental change, virtually independent of its selection coefficient. Second, establishment of the rescue type after the environmental change is hampered unless the wildtype is depleted fast enough.

### The rate of sex

The role of segregation in a diploid one-locus model, where individuals reproduce sexually with probability *σ*_sex_, is very similar to the role of recombination in a haploid two-locus model (see SI section S3 and Otto (2003)), and we can transfer methods and results obtained for the role of recombination in Uecker and Hermisson (2016) to the current problem (cf. compare also Figures 1 and 2 in the present article to Figures 3 and 4 in Uecker and Hermisson (2016)). As in Uecker and Hermisson (2016), we focus on large populations in which we can describe the number of wildtype homoyzgotes and heterozygotes deterministically, both before and after the environmental change. The mutant homozygote, however, will usually be rare initially such that its dynamics requires a stochastic treatment. In order to assess the stochasticity in the number of mutant homozygotes, we apply branching process theory. For details on the analysis, we refer to SI sections S2 and S3 and to Uecker and Hermisson (2016).

#### The role of the dominance coefficient

As known from classical population genetics, whether segregation in sexual reproduction speeds up or slows down adaptation strongly depends on the fitness relations between genotypes. For models in continuous time, the critical factor is the dominance coefficient on an additive scale. For models in discrete time (as considered in this article), dominance on a multiplicative scale is more relevant (Chasnov, 2000; Otto, 2003). Sexual reproduction speeds adaptation up if fitnesses across alleles at one locus are submultiplicative and slows adaptation down if fitnesses are supermultiplicative (Chasnov, 2000; Otto, 2003). (A multiplicative fitness scheme corresponds to a dominance coefficient 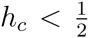 on an additive scale.) In models of evolutionary rescue, the population faces two different environments with a potential shift in the dominance coefficient between environments (Gerstein et al., 2014). In order to quantify deviations from multiplicativity in both phases, we define

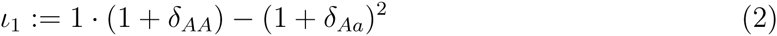

for the old environment and

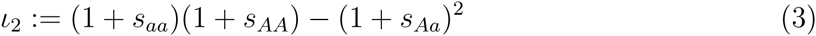

for the new environment. Comparing to a haploid two-locus model, the measure *ι* corresponds to epistasis and takes an analogous role (see SI section S3 and Otto (2003)). Just as epistasis generates linkage disequilibrium, non-zero *ι* leads to an under-or overrepresentation of homozygotes as measured by the inbreeding coefficient

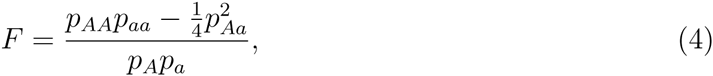

where *p*_*aa*_, *p*_*aa*_, and *p*_*aa*_ denote the relative frequencies of the three genotypes and *p*_*A*_ and *p*_*A*_ the relative frequencies of the two alleles. This deviation from Hardy-Weinberg equilibrium is broken apart by segregation (cf. recombination counteracting linkage disequilibrium). For *ι <* 0, segregation increases the number of homozygotes (speeding adaptation up), whereas for *ι >* 0, it decreases them (slowing adaptation down). For the time after the environmental change, we consider two extreme scenarios, which are examples of *ι*_2_ *<* 0 and *ι*_2_ *>* 0, respectively. In scenario (1), the wildtype is lethal, i.e. *s*_*AA*_ = −1. In scenario (2), the mutant allele is recessive or underdominant, i.e. *s*_*aa*_ ≥ *s*_*Aa*_. We combine these two scenarios with either *ι*_1_ *<* 0 or *ι*_1_ *>* 0 before the environmental change.

In the original environment, mutant homozygotes get generated at approximately constant rate 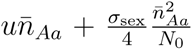, where 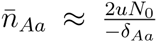 is the expected number of heterozygotes in the standing genetic variation. By mating with the wildtype homozygote, they are broken up at rate ∼ *σ*_sex_, leading to a reduced “effective fitness” *δ_AA_ − σ*_sex_. With these ingredients, we can derive the distribution of mutant homozygotes with the help of a branching process with immigration (where immigration corresponds to the generation of mutant homozygotes through mutation and segregation). The mean number of *AA* mutants in the standing genetic variation can be approximated by 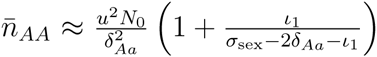 (see SI section S3; the approximation assumes weak selection and *δ_AA_* ≈ 2*δ*_*Aa*_ + *ι*_1_). If the number of mutant homozygotes is large enough to ignore fluctuations around its mean, the probability of rescue from mutant homozygotes from the standing genetic variation can be approximated by

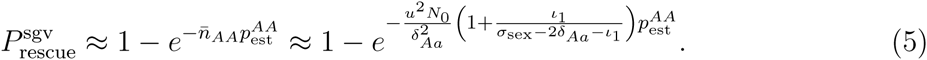

If the wildtype is lethal in the new environment, 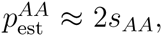, independently of the rate of sex. We can then read off from Eq. (5) that the probability of rescue increases with the rate of sex if *ι*_1_ *<* 0 and decreases with the rate of sex if *ι*_1_ *>* 0 as expected from the classical theory outlined above. Note also how the importance of *ι*_1_ diminishes as the level of sex increases.

In scenario (1), with *s*_*aa*_ = −1 (lethal wildtype, *ι*_2_ *<* 0), the population dynamics after the environmental shift is – at least initially – determined by the heterozygotes, *N* (*t*) *≈n*_*Aa*_(*t*). Mutant homozygotes arise at rate 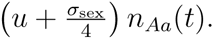. Since one half of all matings between heterozygotes leads to a homozygote offspring, the number of heterozygotes decays at rate 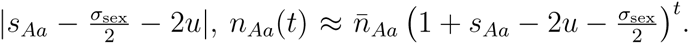. Until extinction, they hence produce 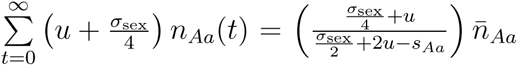 mutant homozygotes. Since the wildtype is lethal, the mutant homozygotes cannot be broken up anymore in the new environment; once generated, they establish with probability 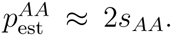. Consequently, rescue from de-novo generated rescue individuals is given by

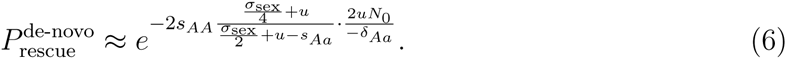

Figure 1 combines this result with *ι*_1_ *<* 0 (Panel A) and *ι*_1_ *>* 0 (Panel B) in the original environment. For *ι*_1_ *<* 0, the effect of segregation is positive in both environments, and rescue increases with the rate of sex. For *ι*_1_ *>* 0, in contrast, the effect of segregation changes from negative to positive upon the environmental change (recall that *ι*_2_ *<* 0). As a consequence, *P*_rescue_(*σ*_sex_) displays an intermediate minimum.

**Figure 1:**
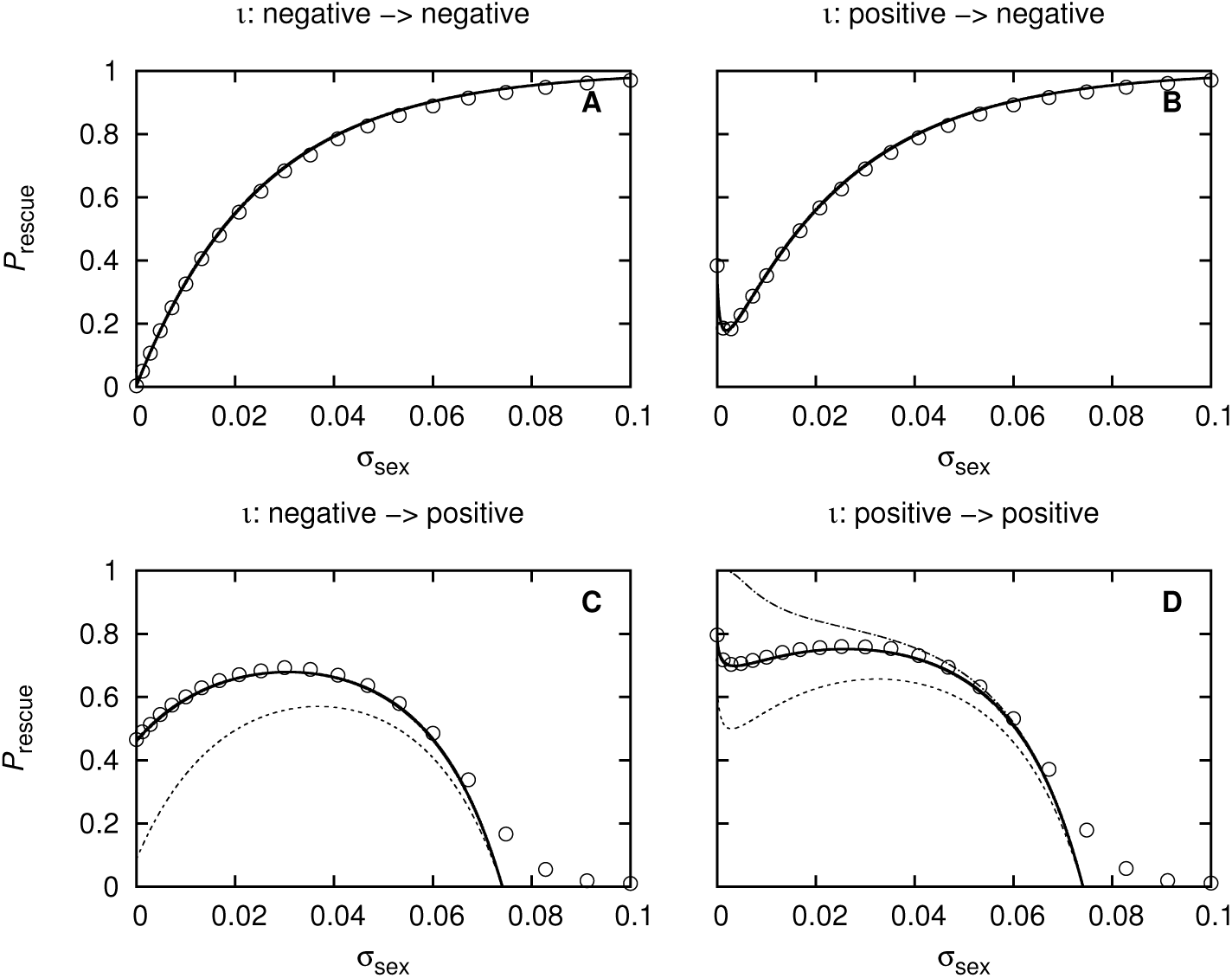
Probability of evolutionary rescue as a function of the probability of sex for different fitness schemes before and after the environmental change. The parameter *ι* measures dominance at a multiplicative scale. For *ι <* 0, sex speeds adaptation up; for *ι >* 0, it slows adaptation down. As the environment shifts, *ι* might change sign, reversing the effect of sex. As examples for *ι <* 0 and *ι >* 0 in the new environment, we consider the special scenarios *s*_*aa*_ = −1 (lethal wildtype) and *s*_*aa*_ = *s*_*Aa*_ (recessive mutant allele). Solid line: total probability of rescue; dashed line: probability of rescue from the standing genetic variation (i.e., without new mutations after the environmental change). In Panels A and B, both lines coincide. Dashed-dotted line in Panel D: probability of evolutionary rescue, ignoring stochastic fluctuations in the number of *AA* mutants before the environmental change. For the analytical predictions, see SI section S3 and Uecker and Hermisson (2016). Parameter values: *N*_0_ = 10^8^, *u* = 2 • 10^−6^, *δ*_*AA*_ = −0.01; first row (A+B): *s*_*Aa*_ = −0.5, *s*_*Aa*_ = −1, *s*_*Aa*_ = 0.002, and *δ*_*AA*_ = −0.1 (i.e., *ι*_2_ *≈ −*0.08, Panel A), *δ*_*AA*_ = −0.0001 (i.e., *ι*_1_ *≈* 0.02, Panel B); second row (C+D): *s*_*Aa*_ = *s*_*AA*_ = −0.03, *s*_*AA*_ = 0.08, and *δ*_*AA*_ = −0.1 (i.e., *ι*_1_ *≈ −*0.08, Panel C), *δ*_*AA*_ = −0.0001 (i.e., *ι*_1_ *≈* 0.02, Panel D). Symbols denote simulation results. Each simulation point is the average of 10^5^ replicates.

In scenario (2), which is an instance of *ι*_2_ > 0, the wildtype is as least as fit as the heterozygotes (*s*_*aa*_ ≥ *s*_*Aa*_) and the population stays dominated by the wildtype until rescue occurs (if it occurs) and the mutant homozygote takes over. The proportion of heterozygotes changes over time such that the rate at which the rescue type gets generated does not take such a simple form as in the first scenario (but the problem remains analytically tractable). Virtually all matings of the mutant homozygote occur with the wildtype homozygote, breaking it up and reducing its “effective” fitness to 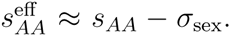. We can then approximate the establishment probability of the rescue genotype by 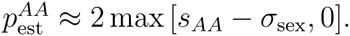. Since *ι*_2_ *>* 0 for scenario (2), the negative effect of segregation is prevalent in the new environment. For *ι*_1_ *<* 0 in the old environment, this means that the role of segregation turns from positive to negative at the time of environmental change, leading to an intermediate maximum in *P*_rescue_(*σ*_sex_); for high rates of sex *σ*_sex_ ≳ *s*_*AA*_, rescue is prevented entirely (Figure 1C). For *ι*_1_ *>* 0 in the old environment, in contrast, based on the fitness scheme, we expect a monotonic decrease of the rescue probability with *σ*_sex_. However, the decrease is not monotonic in Figure 1D. In order to explain this unexpected behavior, we need to consider the effect of stochastic fluctuations in the initial mutant frequency, which we have ignored so far.

#### The role of stochasticity

For simplicity, we consider rescue from the standing genetic variation. Whether rescue occurs or not crucially depends on the number of rescue type individuals that are present in the population at the time of environmental change. This number is a stochastic variable and fluctuates around its mean. For any specific population with a given *n*_*AA*_ at the time of environmental change, the probability of rescue from the standing genetic variation is given by

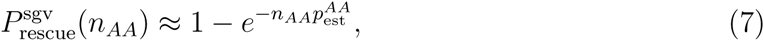

which is a concave function in *n*_*AA*_. This implies that fluctuations of *n*_*AA*_ below the mean 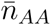 reduce the survival chances of a population more than fluctuations above the mean increase it. Essentially, this follows because a single successful mutant that establishes a permanent lineage is sufficient for population rescue; a second successful mutant does not help (the population would simply be “rescued twice”), while the population goes extinct if no mutant is successful. Overall (i.e. after averaging over the distribution of *n*_*AA*_), stochastic fluctuations in the number of mutant homozygotes therefore diminish the probability of evolutionary rescue. By counteracting any deviations from Hardy-Weinberg equilibrium, segregation dampens fluctuations in the number of mutant homozygotes and hence mitigates this negative effect of stochasticity. The dotted line in Figure 1 shows the probability of evolutionary rescue for the parameters of Panel D if we ignore fluctuations in *n*_*AA*_ prior to the environmental change (hence applying Eq. (5)). For low levels of sex, stochastic fluctuations reduce *P*_rescue_ significantly; with increasing rate of sex, the difference between the curves with and without stochasticity diminish. A similar reasoning holds for de-novo generated mutant homozygotes. The effect is particularly pronounced when the population size has declined to small numbers and stochastic fluctuations in all genotypes are strong. In particular, presence of the rescue type in sexually reproducing species right before the wildtype goes extinct and cannot break the mutant homozygote up anymore increases the probability of rescue considerably. Sexually reproducing populations can therefore have an advantage despite of *ι*_2_ being significantly larger than 0.

#### The population dynamics

From the foregoing discussion, it is apparent that the wildtype takes a double role in the process. On one hand, wildtype individuals can have mutant offspring, providing the raw material for rescue. On the other hand, mutant homozygotes are broken up if they mate with wildtype individuals. Presence of the wildtype after the environmental change has hence antagonistic effects on rescue.

For clonal populations, persistence of the wildtype has no negative consequence, and *P*_rescue_ hence increases with the wildtype fitness 1 + *s*_*aa*_ (solid line in Figure 2A). For a fully sexual population, in contrast, the mutant homozygote is broken up at a very high rate if the wildtype is common. This can result in a decrease of *P*_rescue_ with the wildtype fitness (dotted line). Thus, counterintuitively, the chances of population rescue can be higher if the wildtype gets depleted fast. In the Figure, for intermediate rates of sex, the probability of rescue is non-monotonic in the wildtype fitness and displays a minimum at small |*s*_*aa*_| (dashed lines). Note that sexual reproduction turns beneficial for a value of *s*_*aa*_ that corresponds to a supermultiplicative fitness scheme. This is due to stochastic effects as discussed in the pervious paragraph.

**Figure 2:**
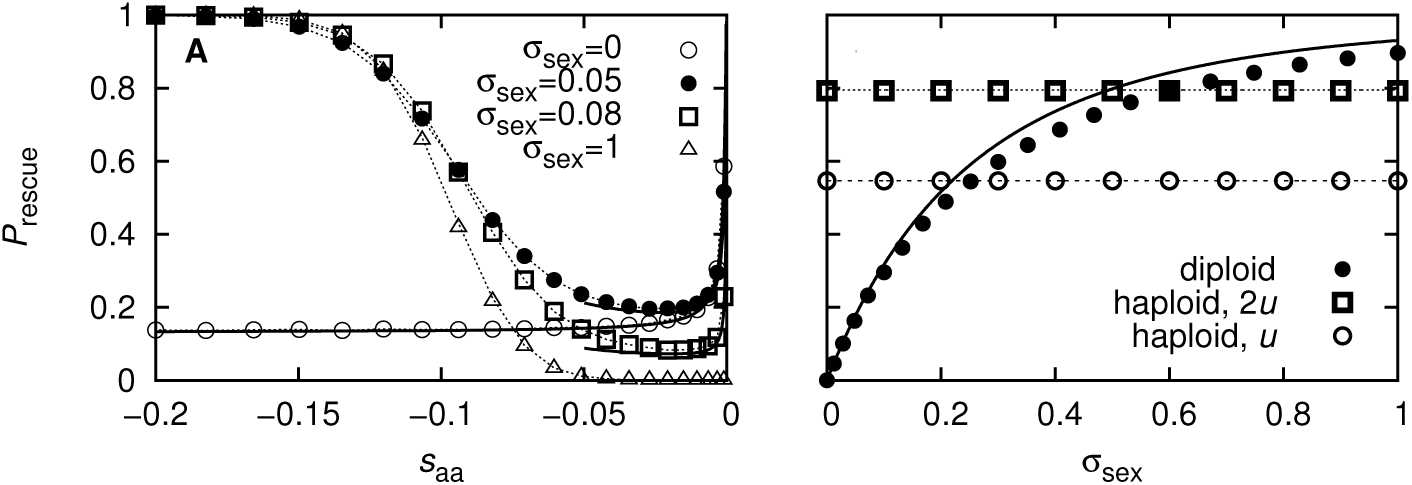
Probability of evolutionary rescue. Panel A: Probability of evolutionary rescue as a function of wildtype fitness for various values of *σ*_sex_. Solid curves consitute analytical predictions (see Eq. (S5.9) with *r* = 0 for *σ*_sex_ = 0 and SI section S3 for *σ*_sex_ *>* 0). Dotted lines interpolate between simulation points and are included to guide the eye. Parameter values: *δ*_*AA*_ = −0.01, *δ*_*AA*_ = −0.0199, *s*_*Aa*_ = −0.05, *u* = 10^−5^, *N*_0_ = 10^6^, *s*_*Aa*_ = 0.1. Panel B: Probability of evolutionary rescue in a diploid sexually reproducing population and in a haploid clonally reproducting population. The solid line shows rescue in a population of diploid individuals with mutation probability *u*; the dashed line shows rescue in a haploid population with mutation probability *u*; the dotted lines shows rescue in a haploid population with mutation probability 2*u*. For the analytical predictions, see SI section S3 (for diploid populations) and Eq. (S2.2) (for haploid populations). Parameter values: *u* = 2 • 10^−6^, *N*_0_ = 10^7^, *δ*_*Aa*_ = −0.01, *δ*_*AA*_ = −0.1, *s*_*aa*_ = −1, *s*_*Aa*_ = −0.5, *s*_*AA*_ = 0.002. Symbols denote simulation results. Each simulation point is the average of 10^5^ replicates.

#### Comparing to a clonal haploid population

The crucial disadvantage of diploid clonal populations consists in the inefficient generation of the double mutant. How does adaptation in a haploid clonal population compare to adaptation in a diploid sexual population, when a single mutant in the haploid population has the same fitness as a mutant homozygote in the diploid population? In the haploid population, only one mutational step is required for adaptation to the new conditions, making rescue a priori seemingly easier. However, as Figure 2 demonstrates, it is nevertheless not necessarily more likely to occur than in sexual populations. Figure 2B considers a scenario with *s*_*aa*_ = −1 (cf. Eq. 5 and Eq. 6). In the clonal population, adaptation hence relies entirely on the rescue mutants in the standing genetic variation, which are present at absolute frequency 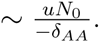 In (partially) sexual populations, −*δ*_*AA*_ mutant homozygotes can still be generated after the environmental change via mutation or mating of mutant heterozygotes. At the same time, due to the lethality of type *aa*, sexual reproduction has no negative effect on the establishment of the mutant homozygote. Figure 2B shows that then, overall, *P*_rescue_ can be higher in partially sexual populations, where mutant heterozygotes buffer the environmental change, than in a haploid clonal population (actually, even the standing genetic variation can contain more mutant homozygotes than in a haploid population; not shown). Even if we double the mutation probability in the haploid population in order to compensate for the smaller mutational target size in haploids, rescue remains slightly more likely in sexual populations (see the dotted line in Figure 2B).

#### Clonal population with mitotic recombination

Up to now, we assumed that in diploid clonal populations, the rescue type can only be created by two-step mutation, which is rather inefficient. However, mitotic recombination provides organisms with an alternative way to reach homoyzgosity. This holds true both for unicellular organisms and for multicellular organisms, when mitotic recombination happens in the germline. Mitotic recombination has been shown to speed up adaptation tremendously in populations of constant size (Mandegar and Otto, 2007; Gerstein et al., 2014). We denote by *r* the probability at which heterozygous mother cells produce homozygous daughter cells through mitotic recombination. Half of these daughter cells are mutant homozygotes. The rescue genotype gets hence generated at rate 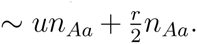. Mathematically (but not biologically), this is very similar to a clonal population with a little bit of selfing. To give an example, Mandegar and Otto (2007) estimate the average rate of mitotic recombination in *Saccharomyces cerevisiae* to be approximately 0.8 × 10^−4^ per cell per generation.

Figure 3 shows that mitotic recombination significantly enhances rescue in clonal populations. However, only for high rates of recombination (relative to estimates in *S. cerevisiae*) does it get more likely than in sexual populations.

**Figure 3:**
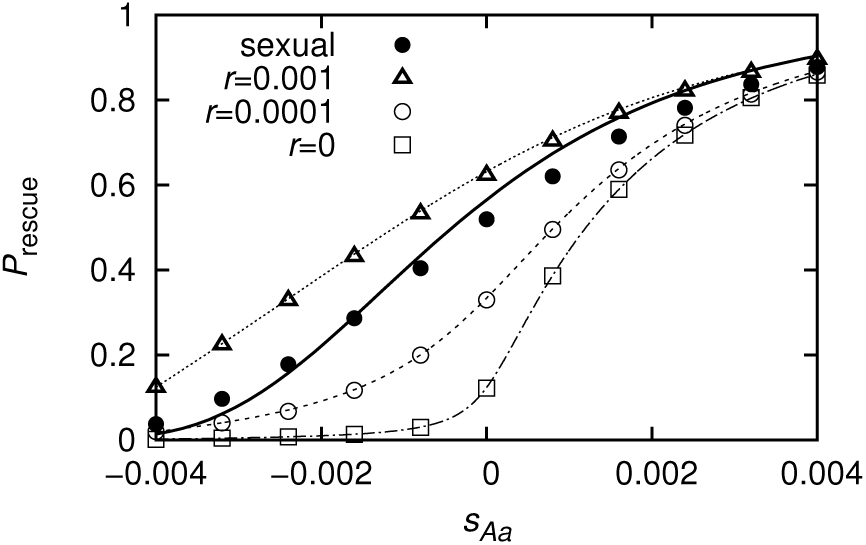
Probability of evolutionary rescue in the presence of mitotic recombination. The figure compares rescue in a sexually reproducing population with clonal populations with different degrees of mitotic recombination. Note that the relative performance of sexual and clonal populations is largely dependent on the impact on low frequency genotypes (“stochastic effect”) and not on the form of selection (multiplicative or not). Panel B: The figure shows by how much mitotic recombination increases the chance of rescue for various values of *s*_*Aa*_. The analytical predictions are based on Eq. (S5.9) and Eq. (S6.26). The expression for *P*_rescue_ in Eq. (S6.26) got evaluated for a discrete set of points that were connected to give a continuous line in the figure. Parameter values: *δ*_*AA*_ = −0.01, *δ*_*Aa*_ = −0.005, *s*_*aa*_ = − 0.01, *s*_*AA*_ = 2*s*_*Aa*_ −*s*_*AA*_, *u* = 5⋅10^−6^, *N*_0_ = 10^5^. Symbols denote simulation results. Each simulation point is the average of 10^5^ replicates.

### The rate of selfing

In the second part of the article, we explore how selfing affects rescue. For this, we consider a population, where individuals reproduce by selfing with probability *σ*_self_ and by outcrossing (random mating) with probability 1 − *σ*_self_. The mathematical analysis is again based on branching process theory and can be found in SI section S6.

While dominance at a multiplicative scale is critical for whether the probability of rescue decreases or increases with the rate of sex, dominance at the multiplicative scale is not the crucial parameter deciding about the role of selfing. As we see below, it is hard to come up with an equally universal criterion. Dominance is still important but it proves more useful to consider dominance at an additive scale; this is also more in line with previous literature on selection in selfing populations (see e.g. Glémin, 2007; Glémin and Ronfort, 2013). Instead of *ι*_1_ and *ι*_2_, we therefore use the parameters *h*’ and *h* from now on, with the relations *δ*_*Aa*_ = *h*’*δ*_*AA*_ and *s*_*Aa*_ = (1 − *h*)*s*_*aa*_ + *hs*_*AA*_ = *s*_*aa*_ + *h*(*s*_*AA*_ − *s*_*aa*_). Since we assume *δ*_*Aa*_ < 0, it holds *h*’ > 0, and since we assume that the mutant homozygote is the fittest type in the new environment, it holds *h* < 1.

#### The composition of the standing genetic variation and the role of the population dynamics

In a fully selfing population, only half of all offspring of a heterozygote individual are again heterozygotes, reducing the “effective selection coefficient” to 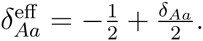 As a consequence, the number of heterozygotes in the standing genetic variation is low with a mean of 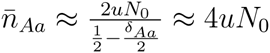 compared to 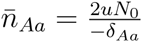 in an outcrossing population. Heterozygotes have offspring of type *AA* with probability 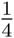 and homozygotes cannot be broken up by segregation, hence 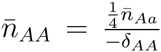 prior to the environmental change. If the mutant homozygote is only weakly deleterious, it accumulates in the population unlike in outcrossing populations where segregation impedes its accumulation. If the mutant homozygote is strongly deleterious, however, both heterozygotes and homozygotes are rare.

Figure 4A compares *P*_rescue_ as a function of *δ*_*AA*_ for a fully selfing and a fully outcrossing population if rescue entirely relies on the standing genetic variation (*s*_*aa*_ = −1). Since type *aa* is lethal in the new environment, the establishment probability of the rescue type is 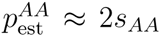 for both populations. In the outcrossing population, in which type *AA* is rare in the standing genetic variation, rescue primarily happens through rescue individuals generated by mating of heterozygotes after the environmental change; rescue is hence basically independent of *δ*_*AA*_. By contrast, in selfing populations, rescue relies on the mutant homozygotes in the standing genetic variation, whose number is strongly affected by *δ*_*AA*_. In Figure 4B, we allow for new mutations after the environmental change (*s*_*aa*_ = −0.01). A slow decay of the wildtype after the environmental switch enhances rescue in a fully selfing population (comparison of Panels A and B). Again, we find that rescue gets completely impeded in the fully outcrossing population (note that *ι*_2_ *>* 0 for the choice of parameters in Figure 4B).

**Figure 4:**
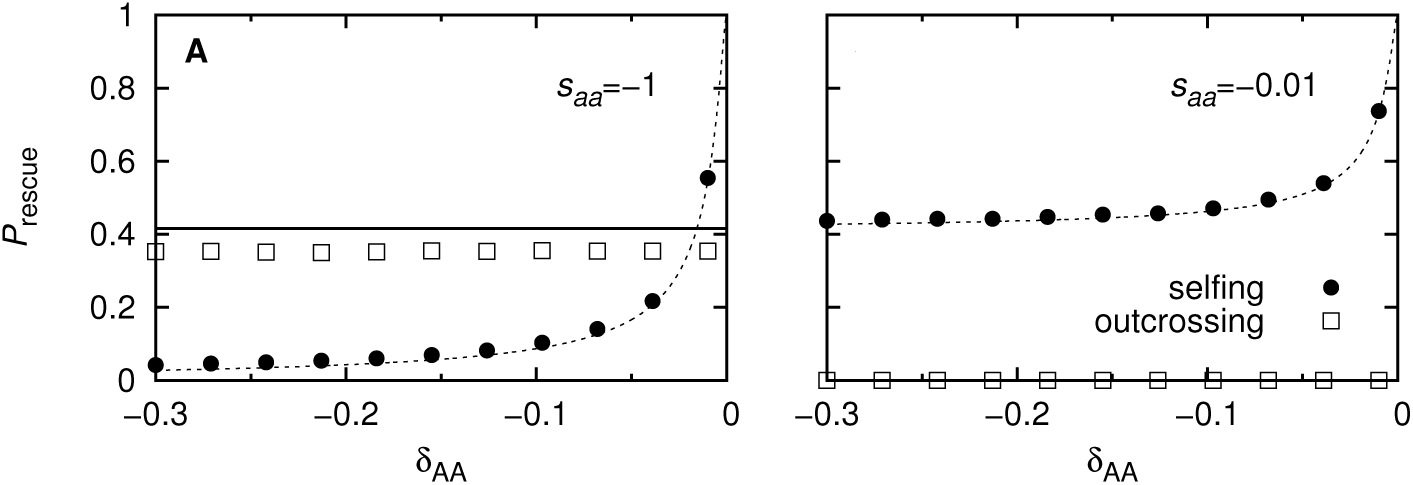
The probability of evolutionary rescue as a function of *δ*_*AA*_ if the wildtype is lethal (Panel A) and if it disappears slowly (Panel B). For selfing populations, the number of *AA* individuals in the standing genetic variation and hence the probability of evolutionary rescue are sensitive to the fitness of type *AA*. In contrast, in randomly mating populations, where type *AA* is continuously broken up, the fitness of type *AA* in the old environment has little influence on rescue. For analytical results, see Eq. (S6.6) and SI section S3. Parameters: *δ*_*Aa*_ = −0.01, *s*_*Aa*_ = −0.3, *s*_*AA*_ = 0.005, *u* = 10^−6^, *N*_0_ = 10^6^. Symbols denote simulation results. Each simulation point is the average of 10^5^ replicates.

A stochastic treatment of rescue in fully selfing populations can be found in SI section S6.1.

#### Effective selection and stochasticity in a selfing population

So far, we focused on the numbers of the three genotypes. We now switch the view and instead consider the number of copies of the mutant allele in the population.

In a selfing population, the mutant allele is very likely to occur in a homozygous individual, which affects the strength of selection it experiences; after the environmental change, it experiences on average stronger selection in selfing than in outcrossing (or clonal) populations. In SI section S6.3.1, we derive an expression for the expected growth of the mutant allele after the environmental change. The analysis is based on a separation of time scales, assuming that genotype frequencies reach equilibrium proportions instantaneously. We obtain

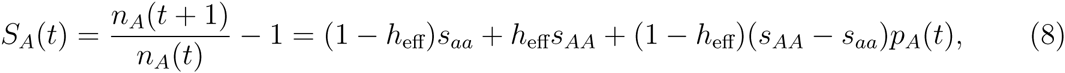

where *p*_*A*_(*t*) is the relative frequency of allele *A* at time *t* and *h*_eff_ = *h* + *F − hF*. We approximate the inbreeding coefficient *F* by its value in the absence of selection and at equilibrium, 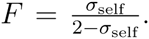 Comparing selfing and outcrossing populations, the difference in growth rate as experienced on average by one copy of the mutant allele is 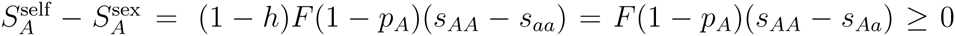 for a given allele frequency *p*_*A*_. Note that 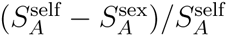 is a decreasing function in *s*_*AA*_. The stronger the effect of the beneficial allele, the less important is the differential growth rate of selfing and outcrossing populations. For the probability of establishment of the mutant allele, not only its expected rate of increase matters but also the strength of stochasticity that it experiences. In SI section S6.3.1, we argue that the variance in the number of descendants of an *A*-allele is given by 1 + *F* (again assuming weak selection and equilibrium) and use this to derive an approximation for the establishment probability of the *A*-allele.

In Figure 5 with 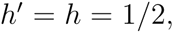 one sees that selfing increases rescue considerably when *s*_*AA*_ is negative and formation of the mutant homozygote is crucial while there is little difference between selfing and outcrossing populations for positive *s*_*AA*_.

**Figure 5:**
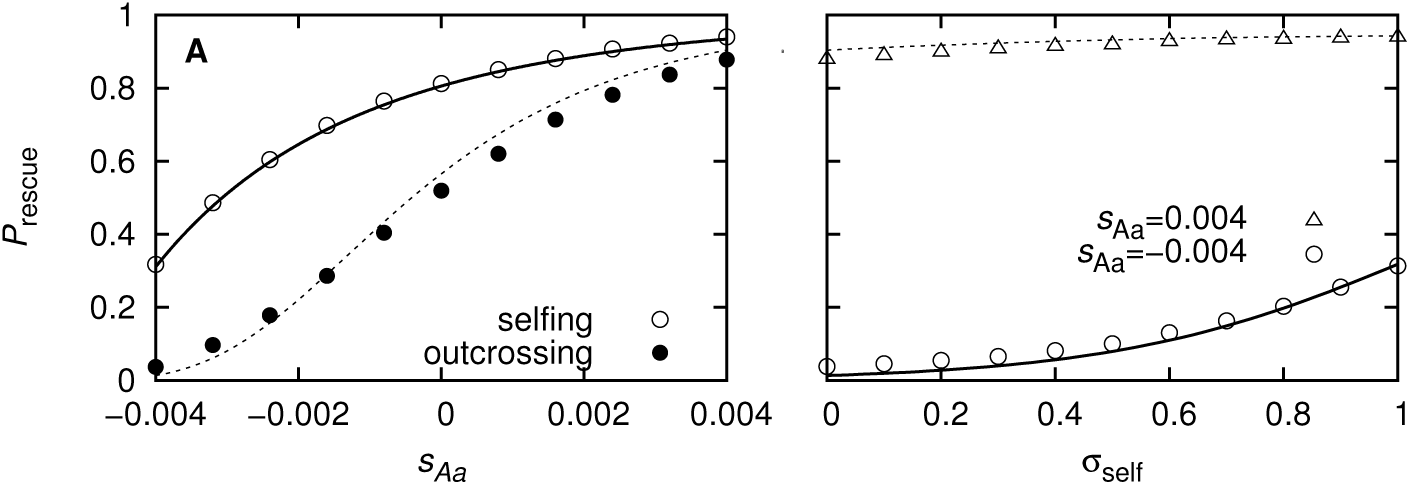
Panel A: Probability of evolutionary rescue for selfing and outcrossing populations as a function of *s*_*Aa*_ when *h* is fixed at 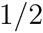 (i.e., the effect of the rescue mutation increases along the *x*-axis). The plot considers the parameter range of negative or only weakly positive *s*_*Aa*_. The analytical approximations are based on Eq. (S6.26) and Eq. (S6.6). Parameter values: 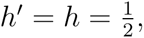 *δ*_*AA*_ = −0.01, *s*_*aa*_ = −0.01, *u* = 5 ⋅ 10^−6^, *N*_0_ = 10^5^. Panel B: Probability of evolutionary rescue as a function of the selfing rate for a weakly and a strongly beneficial mutation. For intermediate dominance 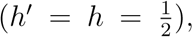 selfing significantly increases the probability of population persistence if the mutation is weakly beneficial (*s*_*Aa*_ < 0), while it has little effect if it is strongly beneficial (*s*_*Aa*_ > 0). The analytical results are based on Eq. (S6.26). Parameter values: *h*’ = *h* = 0.5, *δ*_*AA*_ = −0.01, *s*_*aa*_ = −0.01, *u* = 5 ⋅ 10^−6^, *N*_0_ = 10^5^. The expression for *P*_rescue_ in Eq. (S6.26) got evaluated for a discrete set of p points that were connected to give a continuous line in the figure. Symbols denote simulation results. Each simulation point is the average of 10^5^ replicates.

Note that for *σ*_self_ = 1, the expected growth rate *S*_*A*_(*t*) reduces to *s*_*AA*_ since the allele will (almost) certainly exist in a homozygous individual; it is hence independent of the dominance coefficient. Consequently, the establishment probability of the mutant is independent of *h*, and so is the rate of de-novo generation (2*un*_*AA*_(*t*)) and the frequency in the standing genetic variation (see Eq. (9) below). With this three components, *P*_rescue_ is approximately independent of the dominance coefficient in a fully selfing population.

#### The effect of a change in the dominance coefficient

Finally, we want to elucidate the consequences of a shift in the dominance coefficient for rescue in some more detail. To do so, we first consider/recapitulate the effect of selfing before and after the environmental change.

In section S6.2, we show that the frequency of the mutant prior to the environmental change can be approximated by

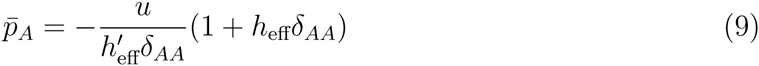

with 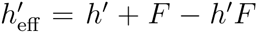 (see also e.g Glémin and Ronfort, 2013); for weak selection (*δ*_*AA*_ small), the result for fully selfing populations derived above coincides with this result. From this approximation, we see that the frequency of the mutant allele in the standing genetic variation decreases with *F* (and hence with the selfing probability) if the dominance coefficient is between zero and one, whilst it increases with *F* if the dominance coefficient is larger than one (i.e., δ_*AA*_ < δ_*AA*_). After the environmental change, the mutation appears by de-novo mutation at rate 2*un*_*AA*_(*t*), independently of the selfing rate. The effect of selfing on the establishment probability is two-fold. Selfing increases the strength of selection on the mutant allele but also the strength of stochasticity that it experiences. For *s*_*Aa*_ < 0, efficient formation of the mutant homozygote is crucial for establishment of the mutant allele, and the positive effect of selfing dominates (this is different if *s*_*Aa*_ > 0 and *h* sufficiently large, see SI section S6.4). Figure 6 shows *P*_rescue_(*σ*_self_) for a switch in the dominance coefficient from *h’* = 0.01 to *h* = 0.5 at the time of environmental change. In this case, the probability of rescue is large in outcrossing populations due to the high initial frequency of the mutant allele. In selfing populations, *P*_rescue_ is large due to the high establishment probability of the mutation. For intermediate selfing, the probability of evolutionary rescue displays a minimum.

**Figure 6:**
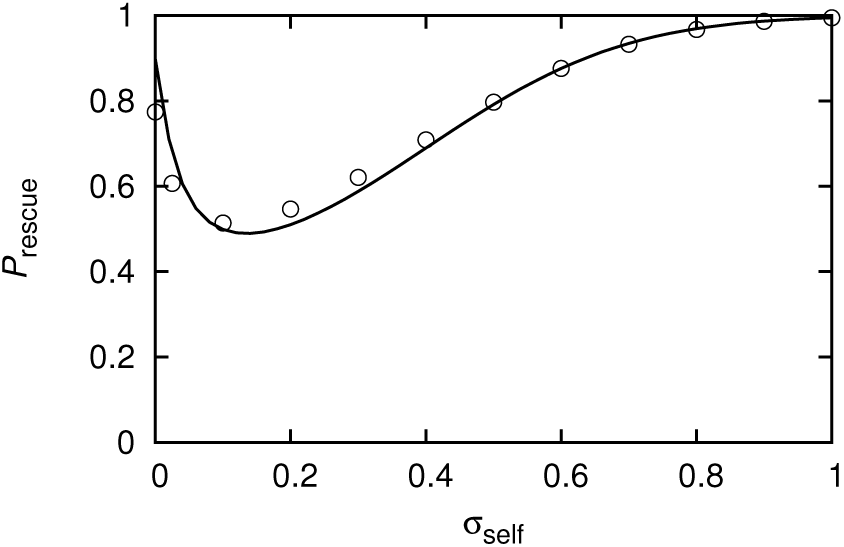
Probability of evolutionary rescue as a function of the selfing probability. The dominance coefficient changes from *h’* = 0.01 to *h* = 0.5 at the time of environmental shift. Outcrossing populations contain the mutant in a high frequency in the standing genetic variation, leading to a high probability of rescue. In selfing population, the initial frequency of the mutation is low but its establishment probability is high due to the efficient generation of the rescue type. The analytical approximation is based on Eq. (S6.26). The expression for *P*_rescue_ in Eq. (S6.26) got evaluated for a discrete set of points that were connected to give a continuous line in the figure. Parameter values: *δ*_*AA*_ = −0.5, *s*_*aa*_ = −0.02, *s*_*AA*_ = 0.01, *u* = 5 ⋅ 10^−6^, *N*_0_ = 10^6^. Symbols denote simulation results. Each simulation point is the average of 10^5^ replicates.

## Discussion

In this article, we investigated the role of the reproductive mode in evolutionary rescue, using a diploid one-locus model. We assumed that the mutant homozygote is the fittest genotype under the perturbed environmental conditions and compared rescue in clonal, selfing, and randomly mating populations.

### The role of the dominance coefficient

Whether random mating increases or decreases the probability of evolutionary rescue compared to selfing or clonal reproduction crucially depends on the dominance coefficient of the mutation before and after the environmental change. Dominance coefficients can range from negative values to values larger than one. Peters et al. (2003) find great variation in the dominance coefficients of mildly deleterious mutations in *C. elegans*. Likewise, Agrawal and Whitlock (2011) observe a broad variety of dominance coefficients, including overand underdominance, for knockout mutations in *S. cerevisiae* (some of them beneficial; they moreover find a correlation between the dominance coefficient and the strength of selection; we ignore such a relationship in this paper). Again for *S. cerevisiae*, Gerstein et al. (2014) show a shift in the dominance coefficient between two different environments with no correlation between the dominance coefficients in the two environments. For our study, we assume that the wildtype is the fittest type before and the mutant homozygote the fittest type after the environmental change but otherwise explore a broad range of dominance coefficients in both phases.

For a comparison of clonal and sexual reproduction, the dominance coefficient (in discrete-time models at a multiplicative scale) is the factor that decides whether sexual reproduction speeds adaptation up or slows it down. A shift in the dominance coefficient upon environmental change can lead to an intermediate minimum or maximum in the probability of rescue as a function of the rate of sex. It is, however, important to note that stochastic effects can be even stronger than the effect of non-multiplicative fitness. By bringing genotype frequencies closer to Hardy-Weinberg equilibrium, sexual reproduction dampens stochastic fluctuations in the relative frequency of mutant homozygotes. In clonal populations these fluctuations decrease the probability of evolutionary rescue.

For selfing vs randomly mating populations, no simple criterion exists, determining whether selfing promotes or impedes adaptation for a given fitness scheme. Glémin and Ronfort (2013) studied the effect of the dominance coefficient on rescue (setting *h’* = *h* in all examples), finding that rescue decreases with the selfing rate for high values of *h* but increases with the selfing rate for low values of *h*. Our work can be seen as complementary to Glémin and Ronfort (2013) in that it explores further parameter regimes and moreover puts an emphasis on the differential role of selfing before and after the environmental change. The mutant frequency in the standing genetic variation decreases with the rate of selfing unless homozygotes have a higher fitness than heterozygotes. For the establishment of the mutant allele after the environmental change in contrast, selfers have an advantage over a broad parameter range due to the efficient formation of the rescue type, which leads to a higher effective growth parameter of the beneficial allele. This advantage increases as the dominance coefficient decreases. The positive effect of stronger selection on the establishment probability is, however, counteracted by stronger stochasticity. If heterozygotes are fit (*s*_*Aa*_ > 0) and the dominance coefficient high, the establishment probability of the mutant allele can therefore indeed be higher in outcrossing than in selfing populations (see Glémin and Ronfort (2013) and SI section S6.4). In the interplay of the potentially opposing effects of selfing on the standing genetic variation and on the establishment probability of the mutant allele in the new environment, the probability of evolutionary rescue can show an intermediate minimum as a function of the selfing rate (see Figures 6 and S6.1). The approximations derived in Glémin and Ronfort (2013) require a high fitness of heterozygotes (at least in populations with a low selfing rate; more precisely *s*_eff_ needs to be significantly larger than zero). In contrast, we include scenarios where the beneficial allele is only slightly beneficial such that the heterozygote fitness is smaller or only slightly larger than one. In that case, establishment of the mutant allele is largely contingent on the formation and persistence of homozygotes. This enlarges the range of dominance coefficients in the new environment for which selfers have a higher probability of rescue than outcrossing species (see also Figure 6 in Glémin and Ronfort (2013)). E.g. if the mutant allele is codominant before and after the environmental shift, *P*_rescue_ is similar for both modes of reproduction if the heterozygote fitness is significantly greater than one in the new environment but selfers have a great advantage if heterozygotes have a low fitness. Note that unlike in randomly mating or clonal populations, the probability of evolutionary rescue in fully selfing populations is insensitive to the dominance coefficient of the mutation for given fitnesses of the homozygote types (see also Figure 5 in Glémin and Ronfort (2013), cf. also Roux and Reboud (2007)). Last, it is important to point out that we do not consider overdominance after the environmental change. In particular if both homozygotes had fitness smaller than 1 and rescue relied entirely on the heterozygote, selfers would suffer a great disadvantage.

### The population dynamics

In clonal or selfing populations, a slow decay of the wildtype population size enhances the probability of population survival since it increases the total mutational input. The same holds true in randomly mating populations with sufficiently fit heterozygotes as studied by Glémin and Ronfort (2013). However, in randomly mating populations with unfit heterozygotes, the breaking up of mutant homozygotes by random mating has a pronounced negative effect on rescue. A fast eradication of the wildtype can therefore promote rescue in randomly mating populations, while a slow decay can hamper it. This observation is of potential relevance for the evolution of resistance in organisms with biparental sexual reproduction such as insects or helminths. The fact that the spread of recessive resistance alleles can be constrained by the presence of wildtype individuals has been brought up multiple times in the resistance literature (e.g. Barnes et al., 1995; Hastings, 2001). In order to apply our theory to the problem of choosing the best dosage (high vs low), it is, however, important to know the dose response curve of all genotypes since most likely, a higher dose not only affects the wildtype (as would correspond to Figure 2A) but also the mutant genotypes.

### Heterozygotes as buffer against environmental change

Heterozygotes, even if they cannot persist longterm in the new environment themselves, can serve as a buffer to environmental change. Consider a situation in which the wildtype is lethal under the new conditions. In randomly mating populations, which usually contain heterozygotes at an appreciable frequency, the rescue type can then be efficiently generated through mating of heterozygotes after the environmental change. Selfing populations, in contrast, harbor few heterozygotes and entirely rely on rescue type individuals in the standing genetic variation. This can put them at a significant disadvantage with respect to outcrossing populations if the rescue type is strongly deleterious in the original environment. Essentially, this corresponds to a situation of small *h’* in the old and large *h* in the new environment.

### The analogy between segregation and recombination

The role of the rate of sex in a diploid one-locus model is analogous to the role of recombination in a haploid two-locus model. The dominance coefficient (at a multiplicative scale for models in discrete time) corresponds to epistasis, and the inbreeding coefficient corresponds to linkage disequilibrium (up to scaling with the allele frequencies). Just as recombination counteracts deviations from linkage equilibrium, segregation counteracts deviations from Hardy-Weinberg equilibrium. As long as the number of heterozygotes/single mutants is large enough to be well described deterministically, results are indeed identical. When random fluctuations in these numbers become important, quantitative differences arise. The probability of rescue is then lower for the haploid two-locus case since both single mutant frequency are subject to stochastic fluctuations. Drift impairs their ratio, making mating between single mutants less likely (see SI section S3 and Figure S3.1). The analogy between the two models is interesting from an experimental point of view. While it is hard to modify the rate of recombination between two loci, it is possible to experimentally control the rate of sex in various organisms such as *S. cerevisiae* or *C. reinhardtii*, making the model predictions experimentally testable.

### Limitations and extensions

We chose the most basic setup to address the problem of how the mode of reproduction influences evolutionary rescue. While this approach allows us to gain a clear picture of elementary processes, it is naturally highly simplifying. First, we only consider one locus. An example for such a simple genetic basis is provided by some cases of insecticide resistance. However, adaptation in natural populations is often polygenic. E.g. successful evolutionary response to climate change normally relies on one or more quantitative traits. With multiple loci, recombination plays an important role. In selfing populations, increased homozygosity (as generated by selfing) alters the action of recombination. Likewise, we neglect background selection which further reduces the variance effective population size in selfing populations (Glémin, 2007).

Second, we assume that the rate of sex and the rate of selfing are constant properties but many species can switch between different modes of reproduction, depending on environmental conditions and the availability of mates. E.g., when stressed, the diploid form of *S. cerevisiae* starts reproducing sexually whereas unstressed, it reproduces clonally by budding (Kassir et al., 1988). Likewise, some plant species reproduce by outcrossing at sunny weather (high availability of pollinators) and by selfing in rainy conditions (Stebbins, 1957). In the simple model used here, the consequences of a switch in the mating system would most likely depend on the dominance coefficients before and after the environmental change. Also, the mode of reproduction is itself subject to selection and can evolve. For example, Bodbyl Roels and Kelly (2011) show that *Mimulus guttatus*, where the selfing rate is a quantitative trait, evolved increased rates of selfing upon pollinator-limitation. Moreover, mating is always guaranteed in our model. However, the mating success of obligate outcrossing individuals often drops with population density since individuals fail to find mates, making rescue less likely to occur. For bi-parental sexual reproduction, we exclusively consider random mating of individuals. Non-random mate choice can considerably influence adaptation. Proulx (1999) show that assortative mating hampers niche expansion in a haploid one-locus two-allele model, while female choice of locally adapted alleles promotes it. Similar effects would probably occur in our model.

We also assume that the mutation rate is independent of the mode of reproduction but Magni and von Borstel (1962) show for *S. cerevisiae* that the mutation rate differs, depending on whether the cells undergo meiosis or mitosis.

Importantly, for bi-parental sexual reproduction, we only consider hermaphrodites and do not allow for two genders/mating types, which might be differentially affected by environmental change. A simple, yet interesting extension of our model would compare rescue relying on a mutation on an autosome vs on the X-chromosome. Urdaneta-Marquez et al. (2014) find that an *X*-linked *dyf-7* haplotype is responsible for resistance against a class of anthelmintics in *Haemonchus contortus*. Based on parameter estimates from this system, they follow the expected genotype frequencies in a classical population genetics model and show that hemizygous resistant males increase the spread of resistance compared to a situation where the allele lies on the autosome.

We also chose the simplest scenario on the ecological side: there is no population structure, and the environmental change (assumed to be sudden) hits the entire population at once. A gradual deterioration of the habitat in space and time alters the population dynamics and along with it the role of segregation and union of gametes in randomly mating populations (see Uecker et al., 2014, for a corresponding haploid model). A very relevant modification of the model would incorporate refugia in which the environment remains benign for the wildtype, leading to its persistence. In a recent experimental study, Lagator et al. (2014) explore the combined effect of sex and migration on adaptation to a sink environment in the algae *C. reinhardtii*. While both sex and migration are beneficial at the initial stage of adaptation, the effect of sex on subsequent adaptation depended on the presence or absence of migration. With ongoing migration, sex slowed further adaptation down by breaking up adaptive gene combinations (since the genetic basis of adaptation did not get determined, it is unclear whether this is due to segregation or recombination). A situation with refugia occurs in agriculture where it is impossible to spray every single leaf with pesticides. Moreover, field margins sometimes remain untreated in order to contain the pesticide safely within the field or to maintain biodiversity and natural enemies of the pest. In both cases, persistence of the wildtype is a side-effect. However, often refugia are preserved with the explicit goal to maintain a wildtype population, which is supposed to hamper the spread of resistance. The idea behind this ‘high-dose-refuge’ strategy is precisely that mutant homozygotes get broken up through mating with the wildtype. Existing models, incorporating this strategy, address for example resistance in insects against transgenic crops that produce *Bt* toxins (Mallet and Porter, 1992; Cerda and Wright, 2004) or herbicide resistance in outcrossing vs selfing weeds (Roux and Reboud, 2007). Beyond these specific models, a generic model along the lines of the present article could shed light on the influence of refugia on the evolution of resistance (see Comins, 1977, for a general, yet mathematically different approach to the problem in obligate sexual insects).

There is no simple answer to the question of which mode of reproduction – clonal reproduction, selfing, or random mating – is best at promoting population survival. The outcome strongly depends on the fitness scheme before and after the environmental change. However, our analysis provides insight into the main principles governing rescue under the three reproductive strategies.

## Acknowledgments

The author thanks Joachim Hermisson, Sally Otto, Nick Barton, Sebastian Bonhoeffer, Sylvain Glémin, Jitka Polechovä, Mato Lagator, Srdjan Sarikas, Richard Gomulkiewicz and two anonymous referees for helpful discussions and/or comments on the manuscript. This work was made possible by a “For Women in Science” fellowship (L’Oréal Österreich in cooperation with the Austrian Commission for UNESCO and the Austrian Academy of Sciences with financial support from the Federal Ministry for Science and Research Austria) and by European Research Council Grants 250152 (to Nick Barton) and 268540 (to Sebastian Bonhoeffer).

## References

Agrawal, A. F. and M. C. Whitlock, 2011. Inferences about the distribution of dominance drawn from yeast gene knockout data. Genetics 187(2):553–566.

Barnes, E. H., R. J. Dobson, and I. A. Barger, 1995. Worm control and anthelmintic resistance: adventures with a model. Parasitology Today 11(2):56–63.

Barrett, S. C. H., 2013. Evolution of Mating Systems: Outcrossing versus Selfing. chap. IV.8, Pp. 356–362, in J. Losos, ed. The Princeton Guide to Evolution. Princeton University Press, Princeton, New Jersey.

Barton, N. H. and B. Charlesworth, 1998. Why sex and recombination? Science 281(5385):1986–1990.

Bodbyl Roels, S. A. and J. K. Kelly, 2011. Rapid evolution caused by pollinator loss in Mimulus Guttatus. Evolution 65(9):2541–2552.

Bourne, E. C., G. Bocedi, J. M. J. Travis, R. J. Pakeman, R. W. Brooker, and K. Schiffers, 2014. Between migration load and evolutionary rescue: dispersal, adaptation and the response of spatially structured populations to environmental change. Preoceedings of the Royal Society B 281(1778).

Carlson, S. M., C. J. Cunningham, and P. A. H. Westley, 2014. Evolutionary rescue in a changing world. Trends in Ecology & Evolution 29(9):521–530.

Cerda, H. and D. J. Wright, 2004. Modeling the spatial and temporal location of refugia to manage resistance in Bt transgenic crops. Agriculture, Ecosystems & Environment 102(2):163–174.

Chasnov, J. R., 2000. Mutation-selection balance, dominance and the maintenance of sex. Genetics 156:1419–1425.

Comins, H. N., 1977. The development of insecticide resistance in the presence of migration. Journal of Theoretical Biology 64:177–197.

Galassi, M., J. Davies, J. Theiler, B. Gough, G. Jungman, P. Alken, M. Booth, and F. Rossi, 2009. GNU Scientific Library Reference Manual. third ed. Network Theory Ltd, Bristol.

Gerstein, A. C., A. Kuzmin, and S. P. Otto, 2014. Loss of heterozygosity facilitates passage through Haldane’s sieve. Nature Communications 5:3819.

Glémin, S., 2007. Mating systems and the efficacy of selection at the molecular level. Genetics 177:905–916.

Glémin, S. and J. Ronfort, 2013. Adaptation and maladaptation in selfing and outcrossing species: new mutations versus standing genetic variation. Evolution 67(1):225–240.

Hartfield, M. and D. Keightley, 2012. Current hypotheses for the evolution of sex and recombination. Integrative Zoology 7:192–209.

Hastings, I. M., 2001. Modelling parasite drug resistance: lessons for management and control strategies. Tropical Medicine and International Health 6(11):883–890.

Kassir, Y., D. Granot, and G. Simchen, 1988. IME 1, a positive regulator gene of meiosis in S. cerevisiae. Cell 52(6):853–862.

Lachapelle, J. and G. Bell, 2012. Evolutionary rescue of sexual and asexual populations in a deteriorating environment. Evolution 66(11):3508–3518.

Lagator, M., A. Morgan, P. Neve, and N. Colegrave, 2014. Role of sex and migration in adaptation to sink environments. Evolution 68(8):2296–2305.

Magni, G. E. and R. C. von Borstel, 1962. Different rates of spontaneous mutation during mitosis and meiosis in yeast. Genetics 47:1097–1108.

Mallet, J. and P. Porter, 1992. Preventing insect adapation to insect-resistant crops: are seed mixtures or refugia the best strategy? Proceedings of the Royal Society of London B 250:165–169.

Mandegar, M. A. and S. P. Otto, 2007. Mitotic recombination counteracts the benefits of genetic segregation. Proceedings of the Royal Society B 274:1301–1307.

Morran, L. T., M. D. Parmenter, and P. C. Phillips, 2009. Mutation load and rapid adaptation favour outcrossing over self-fertilization. Nature 462:350–352.

Otto, S. P., 2003. The advantages of segregation and the evolution of sex. Genetics 164:1099–1118.

Otto, S. P., 2009. The evolutionary enigma of sex. The American Naturalist 174(S1):S1–S14.

Peters, A. D., D. L. Halligan, M. C. Whitlock, and P. D. Keightley, 2003. Dominance and overdominance of mildly deleterious induced mutations for fitness traits in Caenorhabditis elegans. Genetics 165:589–599.

Proulx, S., 1999. Mating systems and the evolution of niche breadth. The American Naturalist 154(1):89–99.

Roux, F. and X. Reboud, 2007. Herbicide resistance dynamics in a spatially heterogeneous environment. Crop Protection 26:335–341.

Schiffers, K., E. C. Bourne, S. Lavergne, W. Thuiller, and J. M. J. Travis, 2013. Limited evolutionary rescue of locally adapted populations facing climate change. Philosophical Transactions of the Royal Society B 268:20120083.

Stebbins, G. L., 1957. Self fertilization and population variability in the higher plants. The American Naturalist 91(861):337–354.

Uecker, H. and J. Hermisson, 2016. The role of recombination in evolutionary rescue. Genetics 202:1–12.

Uecker, H., S. P. Otto, and J. Hermisson, 2014. Evolutionary rescue in structured populations. The American Naturalist 183:E17–E35.

Urdaneta-Marquez, L., S. H. Bae, P. Janukavicius, R. Beech, J. Dent, and R. Prichard, 2014. A dyf-7 haplotype causes sensory neuron defects and is associated with macrocyclic lactone resistance worldwide in the nematode parasite Haemonchus contortus. International Journal for Parasitology 44:1063–1071.

